# Pharmacological activation of AMPK rescues mitophagy defects in models of FBXL4-related mitochondrial DNA depletion syndrome

**DOI:** 10.64898/2025.12.03.692069

**Authors:** Aniketh Bishnu, Robert W. Taylor, Kei Sakamoto, Ian G. Ganley

**Affiliations:** MRC Protein Phosphorylation and Ubiquitylation Unit, School of Life Sciences, University of Dundee, Dundee DD1 5EH, Scotland, U.K; Mitochondrial Research Group, Translational and Clinical Research Institute, Faculty of Medical Sciences, Newcastle University, Newcastle upon Tyne, UK; NHS Highly Specialised Service for Rare Mitochondrial Disorders, Newcastle upon Tyne Hospitals NHS Foundation Trust, Newcastle upon Tyne, NE1 4LP, UK; Novo Nordisk Foundation Center for Basic Metabolic Research, University of Copenhagen, Copenhagen 2200, Denmark

## Abstract

Distinct mitophagy pathways can eliminate not only damaged mitochondria but also healthy ones. In Mitochondrial DNA Depletion Syndrome 13 (MTDPS13), dysregulated BNIP3/NIX-driven mitophagy of functional mitochondria is thought to be the key pathological driver. Patient mutations in the E3 ubiquitin ligase FBXL4 impair the proteasomal degradation of the mitophagy receptors BNIP3 and NIX, causing their accumulation and excessive mitophagy. As a result, mitochondrial content and oxidative phosphorylation decline sharply across multiple tissues, leading to early mortality, with no effective treatments currently existing. In this study, we demonstrate that activating AMPK markedly suppresses BNIP3/NIX-dependent mitophagy and restores mitochondrial respiration in FBXL4-deficient cells. Using both fibroblasts derived from MTDPS13 patients and a chemically-induced in vivo model, we show that small molecule AMPK activation inhibits BNIP3/NIX-mediated mitophagy and recovers mitochondrial content. This work therefore highlights the therapeutic potential of targeting AMPK in MTDPS13.

## INTRODUCTION

Mitophagy refers to several pathways that eliminate mitochondria via the autophagic-lysosomal system (*1, 2*). Traditionally, it is viewed as a quality control mechanism that eliminates damaged mitochondria to help maintain cellular homeostasis. Impaired mitophagy results in the accumulation of dysfunctional mitochondria, a hallmark of several neurodegenerative diseases and other pathologies (*3, 4*). The best-characterized pathway in this context is the Parkinson’s disease–associated PINK1-Parkin cascade, which selectively targets damaged mitochondria for degradation (*5, 6*). However, mitophagy is likely not restricted to damaged/dysfunctional organelles. Activation of the NIX/BNIP3 pathway facilitates mitochondrial turnover in the apparent absence of overt mitochondrial dysfunction, for example during processes such as erythrocyte maturation as well as differentiation of various cell types ranging from neurons to cardiomyocytes (*7–10*). Notably, hyperactivation of NIX/BNIP3-driven mitophagy results in pronounced mitochondrial depletion, as observed in FBXL4-related mitochondrial DNA depletion syndrome (MTDPS13) (*11–13*). This fatal level of organelle loss implies that functional mitochondria are being turned over. In line with this, our recent work has shown mitochondria targeted by NIX-mediated mitophagy maintain functional capacity compared to those degraded by the PINK1-Parkin pathway (*14*).

MTDPS13 (OMIM <u>615471</u>) is a rare human mitochondrial disorder caused by bi-allelic pathogenic variants mutations in the *FBXL4* gene (*15, 16*). FBXL4 (**F-box/LRR-repeat protein 4)** is a mitochondrial outer membrane protein belonging to the F-box protein family and is a component of the SCF (SKP1-CUL1-F-box protein) subfamily of Cullin-RING E3 ubiquitin ligases (CRLs). Within this complex, Cullin 1 provides the structural scaffold, RBX1/2 recruits the E2 ubiquitin-conjugating enzyme, SKP1 functions as the adaptor, and FBXL4 serves as the substrate recognition subunit (*17*). FBXL4 was recently found to interact with partially redundant mitophagy receptors NIX and BNIP3, facilitating their ubiquitination and proteasomal degradation (*12, 13, 18*). This action limits the abundance of NIX and BNIP3, thereby restraining basal mitophagy. In contrast, FBXL4-deficient or mutant cells exhibit elevated NIX and BNIP3 levels, which drive excessive mitophagy and mitochondrial depletion. The resulting loss of mitochondrial mass leads to severe oxidative phosphorylation (OXPHOS) deficiency and broad metabolic disruption (*19*). In vivo, FBXL4 knockout mice demonstrate increased mitophagic flux, reduced mitochondrial content, and perinatal lethality (*20*). Likewise, patients with deleterious, segregating FBXL4 variants typically present with early-onset disease characterized by profound neurodevelopmental delay, encephalomyopathy, hypotonia, and greatly reduced life expectancy (*21, 22*). At present, there are no effective treatments available for FBXL4-associated MTDPS13, although modulation of NIX-dependent mitophagy represents a potential therapeutic avenue.

We recently identified AMPK activation as an inhibitor of NIX-mediated mitophagy (*14*). AMPK is a key regulator of cellular energy homeostasis and a known modulator of autophagy (*23*). It functions as a heterotrimeric kinase complex consisting of a catalytic α subunit (two isoforms), a regulatory β subunit (two isoforms), and a γ subunit (three isoforms). While AMPK activation has classically been associated with the stimulation of autophagy, recent findings indicate that AMPK can also inhibit macroautophagy under certain conditions (*18, 24, 25*). Our previous studies revealed that AMPK phosphorylates ULK1, an autophagy-initiating kinase, promoting its sequestration by 14-3-3 proteins and thereby blocking initiation of NIX/BNIP3-dependent mitophagy in both cell culture models and in mouse tissues. Importantly, AMPK continues to promote clearance of damaged mitochondria through the PINK1-Parkin pathway, independently of ULK1 (*14*).

Given its dual role in suppressing NIX-driven mitophagy and enhancing mitochondrial biogenesis (*23, 26*), we hypothesized that pharmacological activation of AMPK could counteract the pathological overactivation of NIX-mediated mitophagy observed in FBXL4-related MTDPS13. Several direct AMPK activators are under clinical evaluation for conditions including cardiometabolic diseases, metabolic dysfunction–associated steatotic liver disease and steatohepatitis, and polycystic kidney disease (*27–30*). However, whether these compounds can mitigate the excessive mitophagy caused by FBXL4 deficiency has not yet been determined. Accordingly, we aimed to assess whether AMPK activators can restore mitochondrial mass and function by inhibiting NIX/BNIP3-mediated mitophagy in the context of FBXL4-related MTDPS13.

## RESULTS

### Pharmacological activation of AMPK blocks mitophagy induced by FBXL4 inhibition

The levels of NIX and BNIP3 govern the degree of basal mitophagy in cells and are regulated at both the transcriptional level, for example by HIF1α (*9, 31, 32*), and post translational level, via FBXL4-mediated ubiquitylation and proteasomal degradation (*11–13*). As we have previously shown that pharmacological AMPK activation can block NIX/BNIP3 mitophagy induced by HIF1-medaited transcription (*14*), we asked whether AMPK activation could also block NIX/BNIP3-mediated mitophagy caused by inhibition of FBXL4. We have previously demonstrated ARPE-19 cells as a tractable model for evaluating NIX-mediated mitophagy (*33*), and so we treated these with MLN4924, an inhibitor of Cullin Neddylation and hence FBXL4-CRL function (*12, 34*). Upon treatment, and as expected, we observed increased levels of NIX and a concomitant reduction in the mitochondrial proteins HSP60 and COXIV, in both wild-type (WT) and AMPKα1/α2 double knockout (DKO) cells (Fig.1A-B). This indicates FBXL4 inhibition induces NIX-dependent mitophagy in these cell lines. Treatment of cells with MK-8722, a potent and selective pan AMPK activator (*35*), significantly increased AMPK substrate phosphorylation (ACC Ser80 and ULK1 Ser556) in WT but not in DKO cells and had little effect on mitochondrial levels when used alone. In contrast, when used in combination with MLN4924, MK-8722 completely blocked the MLN4924-induced reduction in mitochondrial proteins. This only happened in AMPK WT, but not DKO, cells indicating AMPK activation by MK-8722 blocks NIX-mediated mitophagy in the context of pharmacological FBXL4 inhibition.

**Fig. 1:**
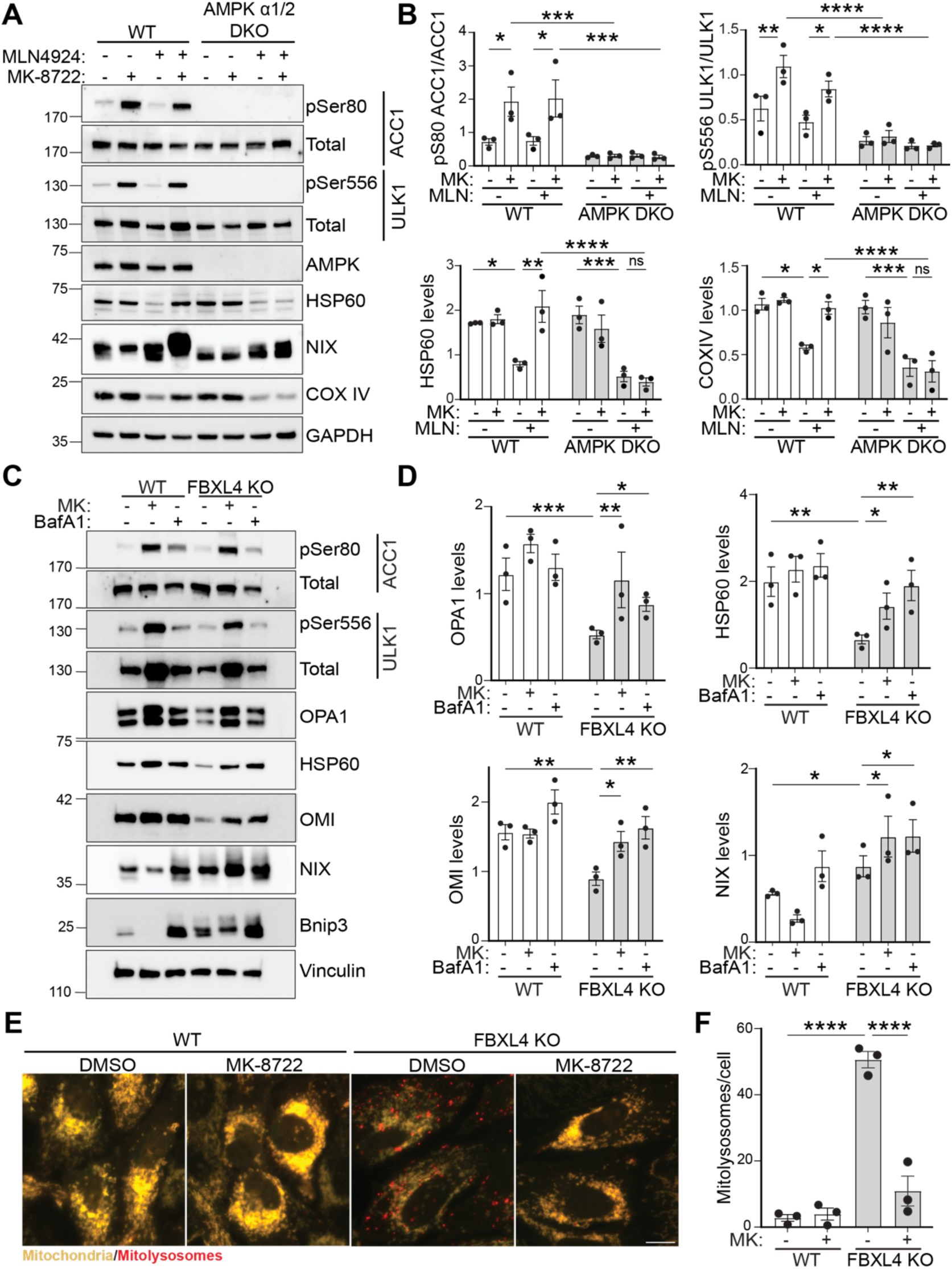
AMPK activation inhibits FBXL4 induced mitophagy. (**A**) Representative immunoblots of indicated proteins from ARPE-19 wildtype (WT) or AMPKα double knockout (AMPK DKO) treated with MLN4924 (5μM) and/or MK-8722 (10μM) for 24 hours. (**B**) Quantification of the immunoblotting data in panel A with protein levels normalised to loading controls. Data represents mean ± SEM (n=3), significance calculated using ordinary two-way ANOVA and Sidak’s multiple comparisons test. (**C**) Representative immunoblots of indicated proteins in ARPE-19 wildtype (WT) and FBXL4 knockout (FBXL4 KO) treated with MK8722 (10μM) for 48 hours and/or Bafilomycin A1(BafA1, 50nM) for final 24 hours. (**D**) Quantification of the immunoblotting data in panel C. Data represents mean ± SEM (n=3), significance calculated using ordinary two-way ANOVA and Tukey’s multiple comparisons test. (**E**) Representative wide-field images of WT and FBXL4 KO ARPE-19 *mito*-QC cells, treated with MK-8722 (10μM) for 48 hours. Red puncta represent mitolysosomes while mitochondrial network appears yellow. Scale bar: 20 μm. (**F**) Quantification of the number of mitolysosomes (red puncta) per cell. Data represents mean ± SEM (n=3), significance calculated using ordinary two-way ANOVA and uncorrected Fisher’s LSD test. For all the experiments statistical significance is depicted as *p<0.05, **p<0.01, ***p<0.001, ****p<0.0001 and ns: nonsignificant.

To specifically evaluate the effect of AMPK activation of FBXL4 induced mitophagy, we generated FBXL4 knockout (KO) ARPE-19 cells using CRISPR-cas9 gene editing. There are currently no validated and commercially available antibodies to detect endogenous FBXL4, yet genomic sequencing revealed the selected clone had a homozygous frameshift mutation leading to the generation of a premature stop at the 183^rd^ codon (See Methods). FBXL4 KO cells displayed increased NIX and BNIP3 levels, along with significant loss of mitochondrial proteins when compared to WT cells (Fig1.C-D). This loss of mitochondrial protein was rescued in the presence of Bafilomycin A1 (BafA1), a lysosomal inhibitor, indicating increased mitophagic flux in the FBXL4 KO cells. Consistent with the MLN4924 experiments, treatment of FBXL4 KO cells with MK-8722 restored mitochondrial proteins to a level comparable to WT cells, showing AMPK activation blocked FBXL4 KO induced mitophagy. Importantly, MK-8722 treatment in FBXL4 KO cells did not decrease NIX levels but rather increased them (comparable to BafA1 treatment), indicating that AMPK activation blocks mitophagy independently of FBXL4-mediated regulation of NIX. To further support the negative effect of pharmacological AMPK activation on mitophagy in FBXL4 KO cells, we also tested two other structurally distinct allosteric AMPK activators: PXL770 (*36*) and BI-9774 (*37*) (Fig.S1A-B). Both compounds activated AMPK, as signified by increased substrate phosphorylation (ACC1 Ser80 and ULK1 Ser556). As with MK-8722, they also significantly increased the level of mitochondrial proteins in FBXL4 KO cells, indicating AMPK-mediated inhibition of mitophagy.

As an alternate way to measure mitophagy, we stably expressed the *mito-*QC reporter in WT and FBXL KO ARPE-19 cells. The reporter works on the principal of lysosomal quenching of GFP fluorescence from a tandem mCherry-GFP tag localized to the outer mitochondrial membrane (*31*). In the cytoplasm, mitochondria fluoresce both red and green, thus appearing yellow/green under the microscope. Upon mitophagy induction, the mitochondria are delivered to the lysosome, resulting in quenching of GFP signal and appearance of red-only punctate structures - indicating mitolysosomes. A significantly higher number of mitolysosomes were observed in FBXL KO cells compared to WT cells, with fewer mitolysosomes observed following MK-8722 treatment (Fig.1E and F). The alternate AMPK activators, PXL-770 and BI-9774, also decreased mitolysosome numbers in FBXL4 KO cells (Fig.S1C-D). Therefore, by two independent measures, we demonstrate that loss of FBXL4 enhances mitophagy, and this can be blocked by AMPK activation.

### AMPK activation restores mitochondrial content and respiration deficiency

Individuals with segregating, damaging FBXL4 variants harbour a significant reduction in mitochondrial content and associated mitochondrial respiratory deficiencies. If the mitochondrial respiration deficit is due to enhanced mitophagy, then blocking mitophagy should restore levels. To evaluate this, we first expressed either the FLAG-tagged WT or a deleterious variant of FBXL4 that harbours a critical mutation in the C-terminal leucine rich repeat region [p.Ile551Asn (*11, 21*)] in FBXL4 KO ARPE-19 cells and assessed the basal and MK-8722-induced level of mitochondrial proteins (Fig.2 A and B). Rescue of cells with WT-FBXL4 significantly reduced NIX levels in comparison to the FBXL4 KO (expressing empty FLAG vector) or the p.Ile551Asn mutant, indicating FBXL4-mediated proteasomal regulation of NIX. Importantly, re-expression of WT, but not p.Ile551Asn mutant FBXL4, resulted in a significantly higher level of mitochondrial proteins OPA1, HSP60, and COXIV, confirming that loss of FBXL4 catalytic activity is responsible for the observed increase in mitophagy. Consistent with previous results, treatment of both FBXL4 KO and p.Ile551Asn mutant cells with MK-8722 restored the level of mitochondrial proteins to basal WT levels. This level was comparable to BafA1 treatment, indicating AMPK activation inhibits mitophagic degradation of mitochondrial proteins. To further confirm the mitophagy levels, we stably expressed *mito*-QC in these cells (Fig2. C-D). As with the immunoblot data, we observed a significant loss of mitolysosomes in FBXL4 KO *mito*-QC cells rescued with WT-FBXL4 compared to cells expressing empty vector or mutant FBXL4. Again, MK-8722 treatment significantly reduced mitolysosome number in both FBXL4 KO and p.Ile551Asn mutant cells. Therefore, MK-8722 treatment blocks NIX/BNIP3-mediated mitophagy and restores mitochondrial content in FBXL4 KO and I551N mutant cells.

**Figure 2:**
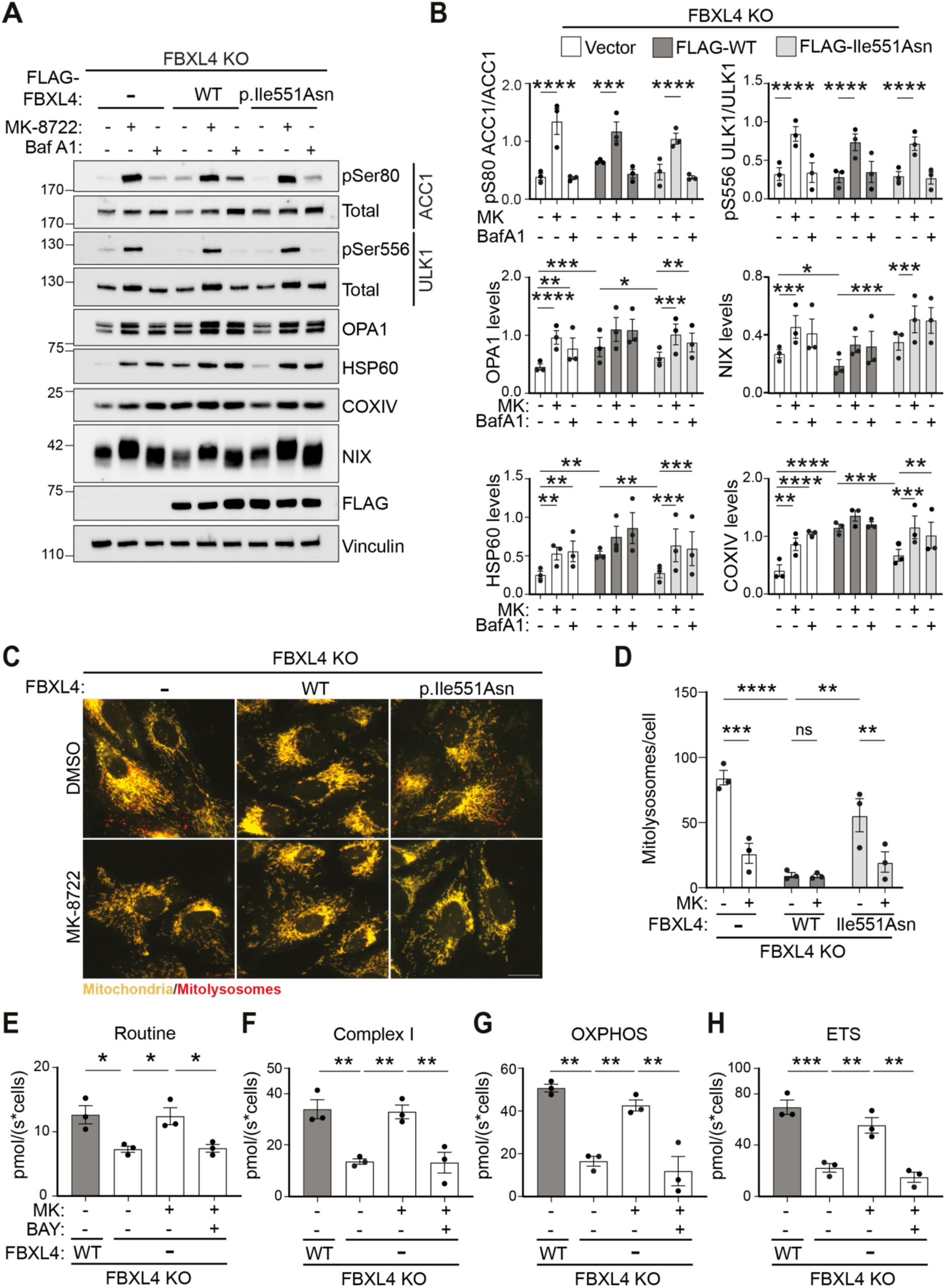
AMPK activation restores mitochondrial content and respiration deficiency. (**A**) Representative immunoblots of indicated proteins from FBXL4 knockout (FBXL4 KO) cells stably expressing either FLAG-empty vector (-), FLAG-FBXL4 wild type (WT) or FLAG-FBXL4 mutant (p.Ile551Asn) treated with MK8722 (10μM) for 48 hours or Bafilomycin A1(Baf A1, 50nM) for final 24 hours. (**B**) Quantification of the immunoblot data in panel A with protein levels normalised to loading controls. Data represents mean ± SEM (n=3), significance calculated using two-way ANOVA and Tukey’s multiple comparisons test. (**C**) Representative wide-field images of FBXL4 KO *mito*-QC cells stably expressing either empty vector (-), FLAG-FBXL4 WT or FLAG-FBXL4 p.Ile551Asn treated with MK8722 (10μM) for 48 hours. Red puncta represent mitolysosomes while mitochondrial network appears yellow. Scale bar: 20 μm. (D) Quantification of the number of mitolysosomes per cell. Data represents mean ± SEM (n=3), significance calculated using two-way ANOVA and Tukey’s multiple comparison test. (**E**) Graph representing routine, Complex I, OXPHOS and ETS respiration measured from FBXL4 KO cells either expressing FLAG-empty vector or FLAG-FBXL4 WT treated with MK-8722 (10μM) or BAY-3827 (5μM) for 48 hours by high-resolution respirometry. Oxygen flux is normalized to the cell number. Data represents mean ± SEM (n=3), significance calculated using ordinary one-way ANOVA and Tukey’s multiple comparison test. For all the experiments statistical significance is depicted as *p<0.05, **p<0.01, ***p<0.001, ****p<0.0001 and ns: nonsignificant.

To confirm that the effects of MK-8722 are AMPK-dependent, we utilized BAY-3827, a potent AMPK-selective inhibitor (*38, 39*) in FBXL4 KO cells (Fig.S2A-B). BAY-3827 alone, or in combination with MK-8722, reduced phospho-ACC1 and phospho-ULK1 levels, confirming AMPK inhibition. The combination treatment also abolished the MK-8722-induced increase in mitochondrial proteins. In line with this, BAY-3827 prevented MK-8722’s ability to block mitolysosome formation (Fig.S2 C-D). This indicates MK-8722 inhibits mitophagy and restores mitochondrial protein content in FBXL4 KO cells via AMPK activation.

Next, we asked whether the mitochondria restored upon AMPK-mediated mitophagy inhibition are functional. We performed high-resolution respirometry in FBXL4 KO cells expressing either empty vector or WT-FBXL4 and measured respiration following stimulation with various mitochondrial complex substrates and specific inhibitors (Fig2.E-H). Cells expressing the WT-FBXL4 had a significantly higher oxygen consumption rate compared to FBXL4 KO cells under all conditions tested: base rate (routine state, Fig.2E), following addition of complex I substrate (CI, Pyruvate-Malate;ADP;Glutamate Fig.2F), complex II substrate (OXPHOS, Succinate Fig.2G) and maximal, uncoupled, rate (ETS, CCCP Fig.2H) as explained in detail in the Materials and Methods. These data demonstrate a significant loss of mitochondrial respiration in FBXL4 KO cells. As AMPK activation can inhibit mitophagy and restore mitochondrial levels in FBXL4 KO cells, we reasoned that this would also restore mitochondrial respiration. Indeed, this proved to be the case as MK-8722 increased oxygen consumption across all conditions to a level comparable with WT cells (Fig.2E-H). This restoration was dependent upon enhanced AMPK activity as it was abolished by co-treatment with BAY-3827. This suggests that the mitochondrial respiratory defects observed in FBXL4 KO cells originate from a loss of mitochondrial number. To confirm this, we measured the flux control ratio and coupling ratio that measure mitochondrial respiration of individual complexes independently of mitochondrial content (Fig.S3A-D). Overall, there were no significant differences between FBXL4 KO cells and those expressing WT FBXL4, indicating that neither FBXL4 KO nor AMPK activation alters the intrinsic mitochondrial oxygen consumption rate under the conditions of the experiment. Taken together, these data suggest that the main consequence of FBXL4 loss is a reduction in mitochondrial quantity rather than quality.

### AMPK activation restores mitophagy levels in patient-derived FBXL4 mutant fibroblasts

To demonstrate disease relevance, we next studied the effect of MK-8722 in patient-derived primary fibroblast cell lines isolated from MTDPS13 patients, utilising cells obtained from patients harbouring homozygous, loss of function truncating *FBXL4* variants. Cells with either the p.Arg435* or p.Gln519* (GenBank RefSeq NM_001278716.2) variants were reported to have significantly diminished levels of mitochondrial DNA and proteins (*15*). In line with these reports, we observed significantly reduced levels of mitochondrial proteins OPA1, OMI, and COXIV, along with a significant increase in NIX and BNIP3 levels, in both the mutant lines compared to WT FBXL4-derived fibroblasts (Fig.3A-B). Importantly, MK-8722 treatment restored the level of mitochondrial proteins to that of WT fibroblasts. Next, to more directly evaluate mitophagy, we stably expressed the *mito*-QC reporter in these cells (Fig.3C-D). A significant increase in mitolysosomes was observed in the FBXL4 mutant fibroblast lines compared to WT cells. Consistent with the immunoblot data, MK-8722 treatment significantly reduced the mitolysosomes numbers back to the WT cell baseline. The data therefore shows that AMPK activation can reduce the level of NIX-mediated mitophagy and restore mitochondrial levels in FBXL4 mutant MTDPS13 patient-derived fibroblasts.

**Figure 3:**
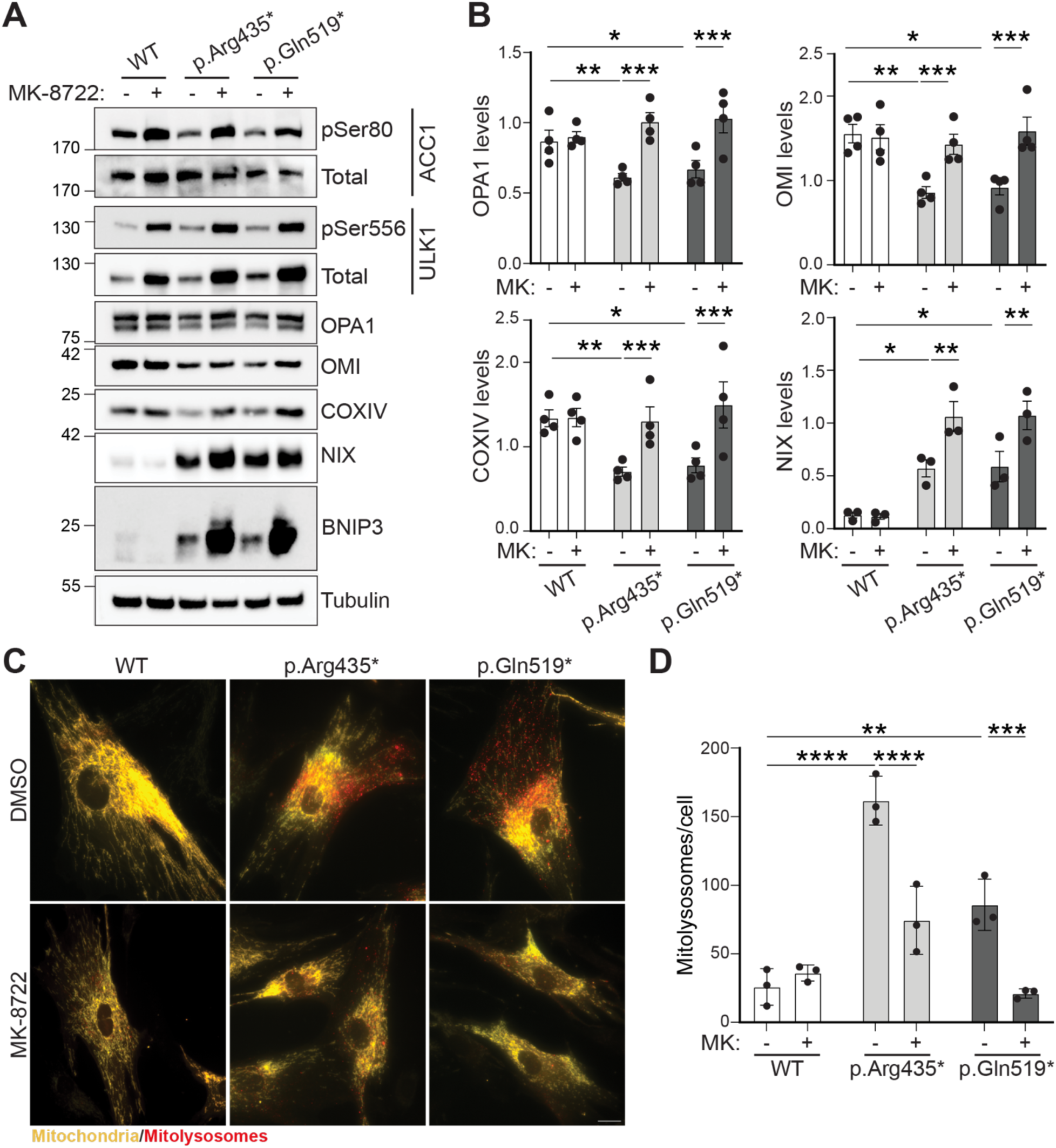
AMPK activation inhibits mitophagy in primary FBXL4 mutant patient fibroblasts. (**A**) Representative immunoblot of indicated proteins in normal (WT) and two FBXL4 mutant, p.Arg435* or p.Gln519*, human MTDPS13 patient fibroblast lines treated with MK-8722 (10 μM) for 48 hours (**B**) Quantification of the immunoblot data in panel A with protein levels normalised to loading controls. Data represents mean ± SEM (n=4), significance calculated using ordinary two-way ANOVA and Tukey’s multiple comparisons test. (**C**) Representative wide-field images of normal WT and p.Arg435* and p.Gln519* FBXL4 mutant human fibroblast lines stably expressing *mito*-QC sensor treated with MK8722 (10 μM) for 48 hours. Red puncta represent mitolysosomes while mitochondrial network appears yellow. Scale bar: 20 μm. (**D**) Quantification of the number of mitolysosomes (red puncta) per cell. Data represents mean ± SEM (n=3), significance calculated using ordinary two-way ANOVA and Sidak’s multiple comparison test. For all the experiments statistical significance is depicted as *p<0.05, **p<0.01, ***p<0.001 and ****p<0.0001.

### AMPK activation reverses mitophagy defects in a chemical-based mouse model of MTDPS13

While we have shown that AMPK activation can rescue FBXL4 mitophagy defects in cell culture models, these systems do not represent true physiological conditions, necessitating further experimental evidence to justify AMPK activation as a potential therapeutic approach to treat MTDPS13. To address this, we used transgenic *mito*-QC mice, which constitutively expresses the *mito*-QC reporter enabling visualisation of mitochondrial architecture and mitophagy within various tissues (*40*). Using this model, we have previously shown that MK-8722 administration reduced basal mitophagy in multiple tissues (*14*), suggesting this represents a viable option for treatment. Unfortunately, constitutive KO of FBXL4 in mice results in perinatal lethality, with most animals dying within 3 days of birth (*11, 20*). Given that conditional FBXL4 KO mice are not yet available, we opted for a pharmacological approach to mimic the disease by inhibiting FBXL4 using MLN4924. Previously, an MLN4924 dose of 60mg/kg in the mouse had been reported to reduce Cullin Neddylation in a human tumour xenograft model (*34*). Thus, we administered the same dose to *mito*-QC mice, either alone or in combination with MK-8722, daily for five consecutive days. Animals were sacrificed at the end of the dosing period on day 5, and tissues were harvested and processed for biochemical and mitophagy assays. AMPK activation results in a sustained lowering of blood glucose levels (*35, 41*) and to confirm this occurred with MK-8722 administration, we measured blood glucose levels pre and post each MK-8722 treatment (Fig.S4A).

As AMPK is a crucial regulator of hepatic metabolic processes and 33% of the reported FBXL4-related MTDPS13 cases show hepatopathy (*42*), we focused on the analysis of liver tissue. Immunoblotting of liver lysates confirmed that MK-8722 administration, either alone or in combination with MLN4924, significantly increased AMPK activity as monitored by phosphorylation of AMPKα at Thr172, ACC1 at Ser79, and ULK1 at Ser555 (Fig.4A-B), which is consistent with the glucose-lowering effect of MK-8722 (Fig.S4A). To evaluate whether MLN4924 administration recapitulates FBXL4-mediated NIX regulation, we monitored the levels of NIX and the mitochondrial proteins ATPB and COXIV. A significant increase in the level of NIX was observed in the MLN4924-treated group compared to the vehicle. Further, MLN4924 alone significantly lowered ATPB levels (with a similar trend for COXIV protein levels), indicating activation of NIX-mediated mitophagy. Co-administration of MK-8722 with MLN4924 significantly impaired the MLN4924-induced reduction in mitochondrial proteins, indicating mitophagy inhibition. To confirm that MLN4924 is indeed inducing mitophagy and that it is impacted by AMPK activation, we analysed the *mito*-QC reporter in liver sections (Fig.4C-D). Firstly, MLN4924 treatment alone significantly increased mitolysosome numbers in the liver, confirming that this compound induces mitophagy. In line with our previous work (*14*), MK-8722 treatment alone reduced liver mitolysosome numbers, demonstrating that AMPK activation reduces overall mitophagy in the liver. Importantly, and consistent with the immunoblot data, MK-8722 administration prevented MLN4924-induced mitolysosome formation in mouse liver.

**Figure 4:**
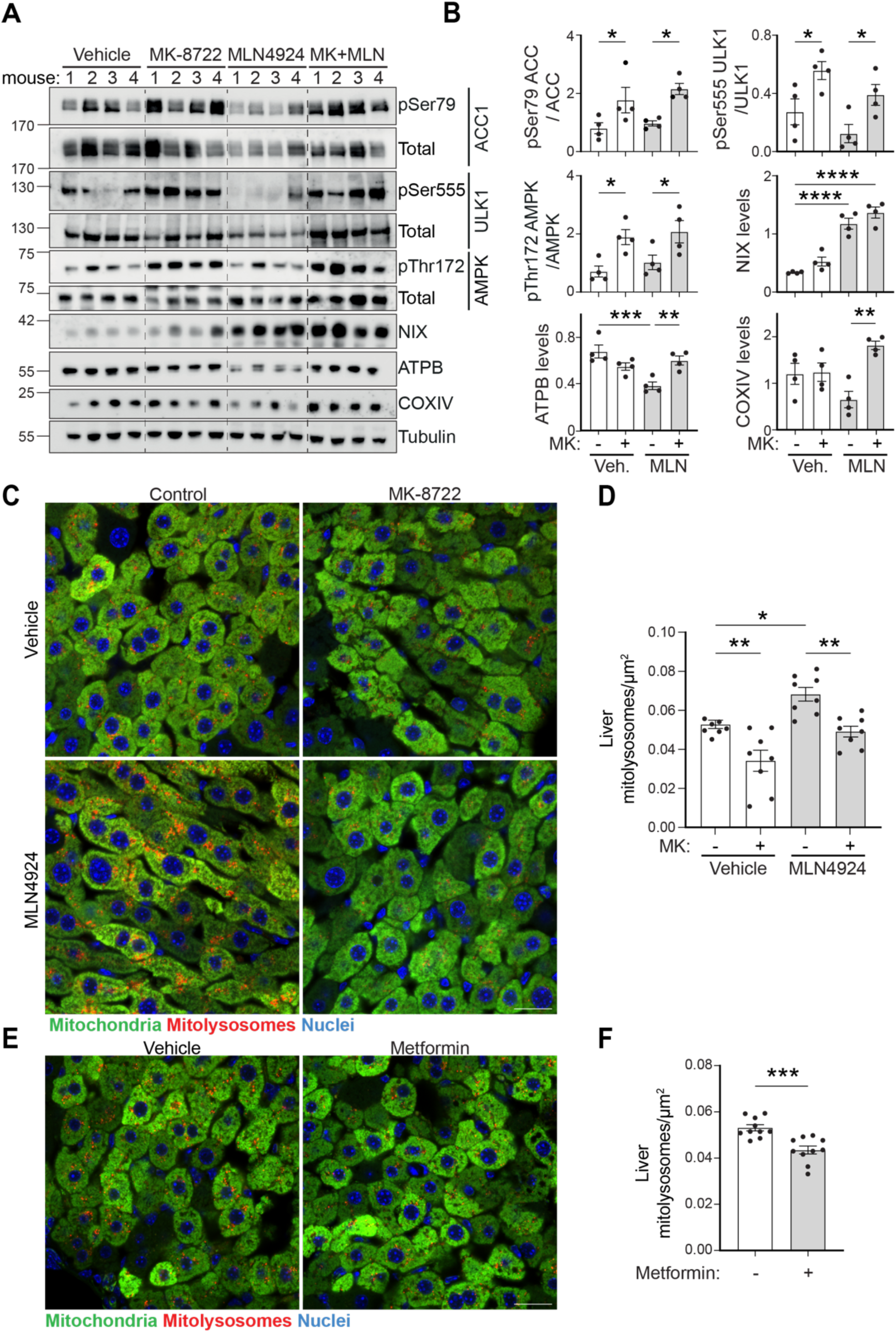
AMPK activation inhibits basal and FBXL4 induced mitophagy in the liver. (**A**) Representative immunoblots of indicated proteins from liver lysates from *mito*-QC mice dosed with vehicle, MK-8722 (30mg/kg) and MLN-4924 (MLN, 50 mg/kg) for 5 days. Each lane represents a different mouse. (**B**) Quantification of the immunoblot data in panel A with protein levels normalised to loading controls. Data represents mean ± SEM (n=4), significance calculated using ordinary one-way ANOVA and Sidak’s multiple comparisons test. (**C**) Representative confocal micrographs of the liver sections from *mito*-QC mice dosed with vehicle, MK-8722 (30 mg/kg) and MLN4924 (MLN, 50 mg/kg) for 5 days. Red puncta indicate mitolysosomes, whereas the mitochondrial network is depicted in green. Scale bar: 20 μm. (**D**) Quantification of the number of mitolysosomes normalized to tissue area. Data represents mean ± SEM (n=8 mice), significance calculated using ordinary one-way ANOVA and Sidak’s multiple comparisons test. (**E**) Representative confocal micrographs of liver sections from *mito*-QC mice dosed with vehicle or metformin (150 mg/kg) for 5 days. Red puncta indicate mitolysosomes, whereas the mitochondrial network is depicted in green. Scale bar: 20 μm. (**F**) Quantification of the number of mitolysosomes normalized to tissue area. Data represents mean ± SEM (n=10), significance calculated using unpaired t-test. For all the experiments statistical significance is depicted as *p<0.05, **p<0.01, ***p<0.001, ****p<0.0001.

Given the effect on liver mitophagy, as well as the therapeutic potential, we also analysed the effects of metformin, the first-line treatment of type 2 diabetes, on liver mitophagy. Since repurposing drugs is both time and cost-effective, and metformin is one of the most prescribed drugs that indirectly activates AMPK (*43*), we reasoned that this analysis could provide valuable insights. *Mito*-QC mice were treated daily with metformin for 5 days at dose of 150mg/kg (*44*), and tissues were then harvested and processed for biochemical and mitophagy assays. Consistent with AMPK activation, metformin administration significantly lowered blood glucose levels (Fig.S4B) and increased substrate phosphorylation (ACC1 at Ser79 and ULK1 at Ser555), as measured by immunoblotting of liver lysates (Fig.S4C-D). To monitor mitophagy, we quantified *mito*-QC in liver sections (Fig.4E-F). Metformin treatment led to a significant reduction in the number of mitolysosomes, indicating metformin-mediated AMPK activation also reduces basal mitophagy in the liver. However, the extent of this reduction was less pronounced than that observed with MK-8722, likely reflecting metformińs lower potency and indirect mechanism of AMPK activation. Taken together, our data show that multiple AMPK activators can modulate liver mitophagy, acting through the BNIP3/NIX-mediated pathway in a negative manner. Considering that metformin primarily targets the liver and exhibits a weaker inhibitory effect on mitophagy compared to MK-8722, we focussed on assessing the effects of MK-8722 in other tissues.

Patients with segregating, pathogenic FBXL4 variants often show severe neurological abnormalities, implying that the homeostasis of NIX/BNIP3-mediated mitophagy in the brain is critical (*42*). We therefore investigated this further in the brains of animals treated with MLN4924 with/without MK-8722. A significant increase in NIX levels was observed in total brain tissue lysate from the animals receiving MLN4924, suggesting inhibition of FBXL4 (Fig.S5A-B). Animals receiving MK-8722, either alone or in combination with MLN4924, showed the expected increase in phosphorylation of AMPK substrates (ACC1 Ser79, ULK1 Ser555) as well as the activating AMPK Thr172 phosphorylation, indicating activation of the AMPK signalling pathway in the brain. We also observed increased NIX levels in the MK-8722 group, which increased even further in combination with MLN4924, indicating AMPK activation may modulate brain mitophagy. To evaluate this, we first focused on the hippocampus, the brain centre responsible for memory consolidation, spatial navigation, and decision-making (*45–47*). We analysed mitophagy in calbindin-positive neurons of the molecular layer of Dentate Gyrus (DG) and Cornu Ammonis (CA1) region (Fig. 5A). MLN4924 significantly increased the number of mitolysosomes in the neurons of the DG compared to the vehicle group, indicating activation of NIX-mediated mitophagy. MK-8722 alone did not alter the number of mitolysosome (Fig5. B-C). However, the combination of MK-8722 with MLN4924 significantly reduced the number of mitolysosome to a level comparable to the vehicle group. An increasing trend in the mitolysosome number following MLN4924 treatment was also observed in the neurons of the CA1 layer, which again was significantly reduced to basal levels in animals co-treated with MK-8722 (Fig5. D-E). Similarly, MK-8722 alone did not alter mitophagy in the neurons of the CA1 layer. This indicates FBXL4 inhibition increases NIX levels and mitophagy in hippocampal neurons, and AMPK activation prevents this. We note, unlike liver, that MK-8722 alone did not alter basal mitophagy in the hippocampal neurons. This could relate to the contribution of different mitophagy pathways within these neurons and the fact that the *mito*-QC assay cannot distinguish between them. We have previously shown that AMPK activation can have contrasting roles in regulating different mitophagy pathways, for example it activates the PINK1/Parkin pathway (*14*), which may be more active in these neurons. Potentially related to this, not all neuronal populations displayed a mitophagy change upon MLN4924 and/or MK-8722 administration. For example, we did not observe any significant changes in the number of mitolysosomes in the Purkinje neurons of the Cerebellum (Fig.S5C-D). This however could also relate to the relative neuron-specific activation of AMPK and further work will be needed to determine this. Importantly though, we show that FBXL4-dysregulated mitophagy is occurring within neuronal populations within the brain and this can be rescued by administration of an AMPK activator. Given that the brain is a major organ affected in MTDPS13, we believe small molecule AMPK activation is a promising therapeutic approach to treat this disorder.

**Figure 5:**
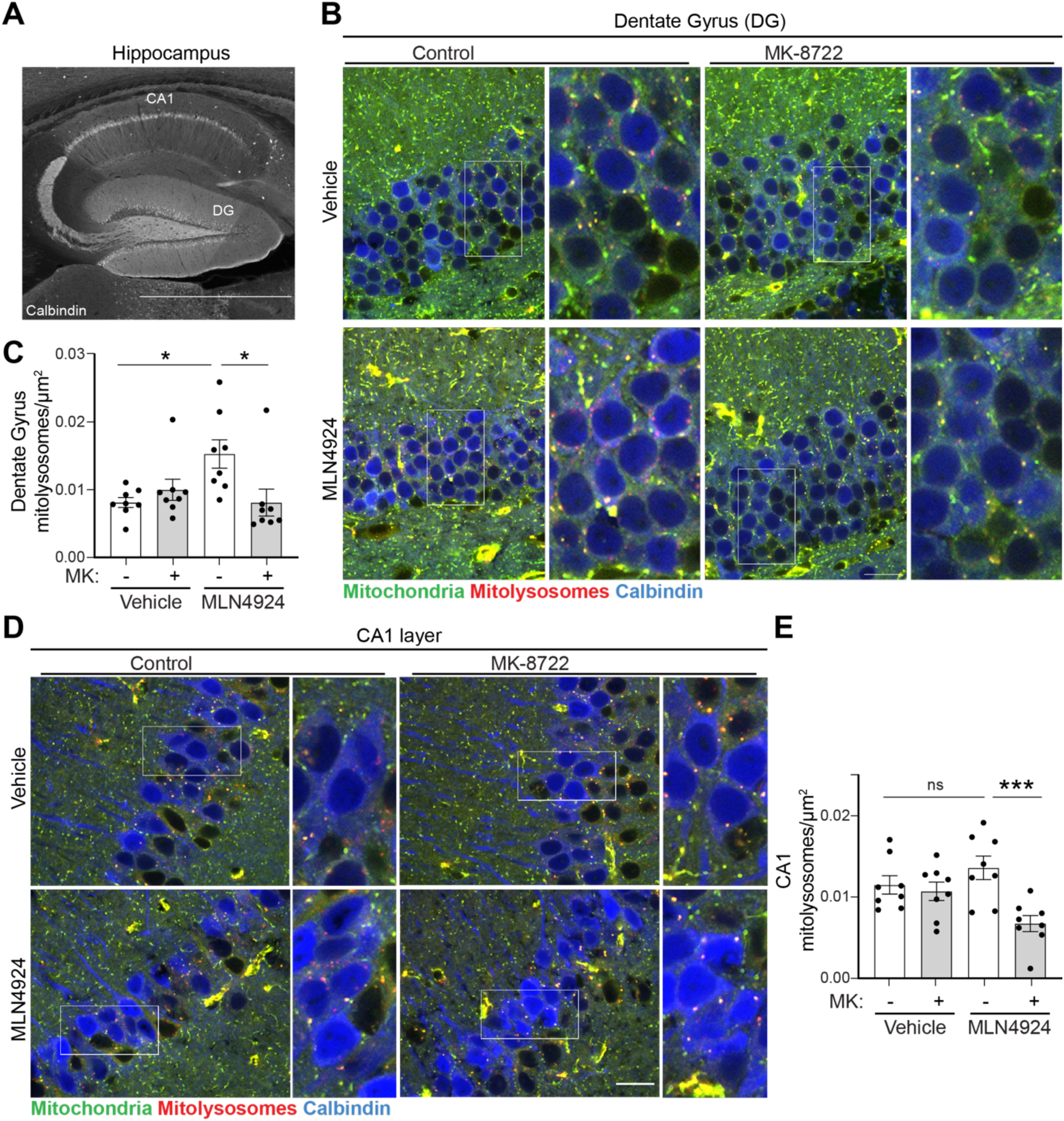
AMPK activation inhibits FBXL4 induced mitophagy in the hippocampus. (**A**) Representative tile scan confocal micrograph of the mouse hippocampus depicting Calbindin positive (Grey) neurons in the molecular layer of Dentate Gyrus (DG) and the Cornu Ammonis (CA1) regions. Scale bar, 5mm. (**B**) Representative confocal micrograph of Calbindin immunolabelled neurons (blue) in the molecular layer of Dentate Gyrus (DG) and (D) Cornu Ammonis (CA) layer from the brain sections of *mito*-QC mice dosed with vehicle, MK-8722 (30 mg/kg) and MLN-4924 (MLN, 50 mg/kg) for 5 days. Enlarged images of the boxed region are shown in the right of respective images. Red puncta indicate mitolysosomes, whereas the mitochondrial network is depicted in green. Scale bar: 20 μm. (**C**) Quantification of the number of mitolysosomes normalized to the area of the calbindin labelled DG neurons. Data represents mean ± SEM (n=8), significance calculated using ordinary one-way ANOVA and Sidak’s multiple comparisons test. (**D** and **E**) as for panels B and C but monitoring calbindin-labelled neurons in the CA1 region. For all the experiments statistical significance is depicted as *p<0.05, ***p<0.001 and ns: nonsignificant.

## DISCUSSION

FBXL4-related MTDPS13 is a rare and often fatal mitochondrial disorder for which no therapeutic interventions currently exist. To date, more than 100 individuals carrying pathogenic *FBXL4* variants have been reported worldwide, supporting detailed clinical phenotyping (*48*). Affected individuals exhibit severe clinical manifestations, including growth retardation, feeding difficulties, hypotonia, and profound developmental delay, with a median age of death of approximately three years (*21*). Disease management sadly remains purely supportive.

Here, we identify pharmacological AMPK activation as a potential therapeutic strategy for MTDPS13. We demonstrate that AMPK activation suppresses excessive NIX/BNIP3-mediated mitophagy, thereby restoring mitochondrial content and improving mitochondrial respiratory function in FBXL4 KO, as well as in patient-derived fibroblasts harbouring FBXL4 loss-of-function truncating variants (Fig.2 and Fig.5). We have previously shown that distinct mitophagy pathways selectively target different mitochondrial subpopulations. Specifically, mitochondria eliminated via the NIX-mediated pathway retain higher membrane polarization and respiratory capacity than those degraded by the PINK1/Parkin pathway following mitochondrial damage (*14*). The current findings further support this preferential degradation of functional mitochondria by NIX-mediated mitophagy as stabilization of NIX in FBXL4 KO cells triggered excessive mitophagy even in the absence of mitochondrial stressors. If the mitochondria targeted by NIX were already severely compromised, their preservation would not have enhanced the mitochondrial respiration levels observed in FBXL4 KO cells (Fig.2E-H). Together, these data underscore the role of NIX-mediated mitophagy in regulating mitochondrial abundance and cellular metabolism. A severe reduction in mitochondrial respiration, caused by uncontrolled NIX/BNIP3-dependent mitophagy, is likely responsible for the energetic and metabolic failure observed in multiple tissues of MTDPS13 patients. Consistent with this model, AMPK activation enhanced mitochondrial respiration in FBXL4 KO cells by restoring mitochondrial content without altering the intrinsic respiratory capacity of individual mitochondria (Fig.S3A–D). Sustained AMPK activation is also known to stimulate mitochondrial biogenesis through PGC-1α (*49, 50*), suggesting that pharmacological activation of AMPK could offer a dual therapeutic benefit in MTDPS13 by simultaneously suppressing excessive mitophagy and promoting mitochondrial biogenesis.

Beyond the context of FBXL4-related mitochondrial disease, our findings refine the prevailing view of AMPK as a universal activator of autophagy and mitophagy. While nutrient deprivation typically activates AMPK to induce autophagic degradation of cellular components for ATP generation, mitochondria themselves are the primary sites of ATP production. Thus, indiscriminate degradation of functional mitochondria would be energetically disadvantageous. We previously demonstrated that AMPK phosphorylates ULK1 at Ser556 and Ser694, facilitating its interaction with 14-3-3 proteins. This interaction sequesters ULK1 and prevents it from phosphorylating downstream targets such as ATG13 and ATG14, thereby inhibiting initiation of NIX-dependent mitophagy (*14*). In agreement with this mechanism, AMPK activation markedly reduced mitolysosome formation in FBXL4 KO/mutant cell lines and patient-derived fibroblasts (Fig. 2C–D and Fig. 5C–D), confirming that AMPK blocks the initiation phase of NIX-mediated mitophagy.

Interestingly, PPTC7, a positive regulator of the FBXL4–SCF complex, is upregulated during starvation, serving to prevent mitophagy activation and protect mitochondria under energy stress (*51*). Proteomic analyses of autophagic cargoes likewise reveal that mitochondria are largely excluded from degradation during nutrient deprivation (*52*). Our findings uncover an additional layer of control by which AMPK prevents the degradation of functional mitochondria via inhibition of NIX/BNIP3-dependent mitophagy. In contrast, AMPK promotes the clearance of damaged mitochondria through the PINK1–Parkin pathway independently of ULK1 (*14*). Together, these observations highlight a dual and context-dependent role for AMPK in mitochondrial quality control—facilitating removal of damaged organelles while preserving healthy ones.

Crucially, we now demonstrate that AMPK activation inhibits excessive mitophagy in an in vivo chemical-based model of MTDPS13 (Fig.3 and Fig.4). Consistent with the neurological features of FBXL4 deficiency, we observed elevated mitophagy in the hippocampus following MLN4924 administration. Activation of AMPK by MK-8722 suppressed MLN4924-induced mitophagy without affecting basal turnover (Fig.4B–E). Although the blood–brain barrier permeability of MK-8722 was not reported (*35, 53*) and remains to be fully established, our findings suggest that systemic AMPK activation is sufficient to modulate neuronal mitophagy.

Drug repurposing offers a rapid and cost-effective route to therapeutic development. Although MK-8722 is a preclinical compound, our findings strongly support the potential of AMPK activators in FBXL4-related MTDPS13. We additionally tested metformin, an indirect AMPK activator widely prescribed for type 2 diabetes and found that it reduced basal mitophagy in the liver (Fig.3E–F). However, metformin’s pharmacokinetic profile may limit its systemic utility: it is primarily absorbed in the liver and intestine via organic cation transporter (OCT) 1 and, to a lesser extent, in the kidney via OCT3 (*43, 54*). Moreover, high doses can cause lactic acidosis, restricting its application in mitochondrial disorders such as MTDPS13 that frequently associate with a metabolic acidosis (*55*).

Of further note, PXL-770, a direct allosteric AMPK activator tested in this study, has advanced to phase IIa clinical trial and demonstrated favourable tolerability in patients with non-alcoholic fatty liver disease (*27*). PXL-770 has been shown to improve mitochondrial function in models of X-linked adrenoleukodystrophy (ALD) and autosomal dominant polycystic kidney disease (ADPKD), and to restore mitochondrial DNA content in kidneys of PKD1 KO (*28, 56*). In our hands, PXL-770 activated AMPK to a comparable extent as MK-8722 (Fig. S1A–B) and was equally effective in suppressing mitophagy and restoring mitochondrial content in FBXL4 KO cells (Fig. S1C–D). These data identify PXL-770 and other clinically applicable small-molecule AMPK activators, as potential candidates for therapeutic repurposing in FBXL4-related MTDPS13.

In summary, our study highlights the inhibitory role of AMPK in the mitophagic degradation of functional mitochondria and provides a mechanistic rationale for targeting AMPK in FBXL4-related MTDPS13. By suppressing NIX-mediated mitophagy and promoting mitochondrial biogenesis, AMPK activation represents a promising strategy to restore mitochondrial homeostasis in this otherwise untreatable disease. These findings warrant clinical evaluation of existing AMPK activators for safety and efficacy, and further development of next-generation direct AMPK activators as potential therapies for mitochondrial DNA depletion syndromes.

## MATERIALS AND METHODS

### 1. Antibodies and Regents

Primary antibodies were used at a dilution of 1:1000 for immunoblotting unless mentioned otherwise: Phospho Ser79 ACC (Cell Signaling Technology (CST) #3661), ACC (CST #3662), Phospho Ser556 ULK1 (CST #5869S), Phospho Ser757 ULK1 (CST #6888S), ULK1 (CST #8054S), OMI (MRC PPU Reagents and Services, University of Dundee, S802C), HSP60 (CST #4870), NIX (CST #12396), OPA1 (BD Bioscience #612606), ATPB (Abcam #ab14730), COX IV (Cat# 4850) AMPK pan-α (CST # 2532S, # 2793S), AMPK pan-β (CST # 4150S), Phospho Thr172 AMPK (CST #2535S), Anti-FLAG (Sigma #F1804) Tubulin (Proteintech #66031-1-Ig, 1:10000), Vinculin (Abcam #ab129002, 1:10000), GAPDH (Proteintech #60004-1-Ig, 1:10000), Calbindin-D28k (Swant, #CB38,1:500).

Secondary antibodies were obtained from Thermo Scientific and used at a dilution of 1:10,000 for immunoblotting: goat anti-Rabbit IgG-HRP (#31460), goat anti-mouse IgG-HRP (#31430), donkey anti-sheep IgG-HRP (#A16041), and Pacific Blue Goat anti-Rabbit IgG (Invitrogen #P10994, 1:200)

Bafilomycin A1 (BML-CM110) was procured from Enzo, MK-8722 (#HY-111363) and PXL-770 (#HY-160004) were brought from MedChemExpress, BI-9774 was kindly provided by Boehringer Ingelheim via its open innovation platform opnMe, available at https://opnme.com, MLN4924 (#5054770001) was purchased from Merck.

### 2. Cell culture, transfection, and infection

ARPE-19 cells and Patient fibroblast lines were cultured in DMEM: F12 media supplemented with 10% Fetal bovine serum, 2mM L-glutamine, 100 U/ml Penicillin, and 0.1 mg/ml Streptomycin. HEK293/FT cells were cultured in DMEM, supplemented with 10% (v/v) FBS, 2 mM L-glutamine, 100 U/ml Penicillin, and 0.1 mg/ml Streptomycin. All cell lines were grown at 37°C with 5% CO_2_ in a humidified incubator. Fibroblasts from affected individuals and age-matched controls were obtained as previously described by informed consent in accordance with the Declaration of Helsinki protocols and local Institutional Review Boards (*15*).

Retroviral transduction was used to generate stable cells expressing the protein of interest. For generating the retrovirus, 70% confluent HEK293FT cells were transfected with 0.5µg of pBABE vector expressing respective genes, 0.33 µg of Gag-Pol, and 0.17 µg of VSVG using X-treme Gene HD (Roche #6366244001) according to the manufacturer’s protocol. The cell supernatant was harvested after 48 hours of transfection and filtered through a 0.45 μm filter to remove floating cells. The filtered supernatant was used to transduce cells at 70% confluency with polybrene (10 μg/ml). After 48 hours of transduction, the cells were selected with either puromycin (2 μg/ml) or hygromycin (100 μg/ml) and the stable pool of cells was used for the experiment.

### 3. Generation of CRISPR cell lines

Two single CRISPR guides, cloned in the pX459 Cas9 vector, were tested individually to generate FBXL4 knock-out cells. Guide 1: GTTTAGCAGTGCTGTCCTCG and Guide 2: GTTGGTCAGAGAGACCTACGA. Guides were transfected in ARPE-19 cells with 6 µl of GeneJuice (Sigma #70967) per µg of DNA. Post 24 hours of transfection, 2 µg/ml of puromycin was added for 24 hours for selecting transfected cells. The cells were then allowed to recover. Single-cell clones were generated by serial dilution and screened using immunoblotting for NIX, as no validated FBXL4 antibody is commercially available. Guide 2 successfully led to increase in NIX levels.

To confirm knockout of FBXL4, genomic DNA was extracted using using Qiagen DNeasy kit following manufacturer’s instruction. The CRISPER cleavage site was PCR amplified using KOD Hot start PCR (Merck #71086) according to manufacturer’s protocol with the following primers (Fwd: GTTGGTTTAAGCCATGCACACAGG, Rev: AGCATGTATTGCC CACAGTTAAAGTG). The PCR product was then cloned into a holding plasmid using the StrataClone Blunt PCR cloning kit (Agilent #240207) and transformed into the StrataClone *E. coli* cells provided in the kit. Plasmids were isolated from multiple bacterial clones using QIAprep Spin Miniprep Kit (Qiagen #27104) following manufacturer’s protocol. Positive clones containing the PCR product were identified with EcoRI digestion (Thermo Fisher, #FD0274). The positive clones were then analysed with sanger sequencing, which revealed a homozygous single-nucleotide deletion causing a frameshift mutation and the introduction of multiple premature stop codons beginning at the 183rd codon.

### 4. Plasmids

pBABE mito-QC retroviral system (DU40799) was used as described previously (*31*). pBABED-Puro-FBXL4-FLAG2 (DU71998), pBABED-Puro-FBXL4-I551N-FLAG2 (DU71999) retroviral constructs were obtained from MRC-PPU reagents and Services.

### 5. Immunoblotting

Cell lysates were prepared using NP-40 lysis buffer (50 mM HEPES pH:7.4, 150 mM NaCl, 1mM EDTA, 10% glycerol, 1% NP-40) supplemented phosphatase inhibitor cocktail (1.15 mM sodium molybdate, 4 mM sodium tartrate dihydrate, 10 mM β-glycerophosphoric acid disodium salt pentahydrate, 1 mM sodium fluoride, 1 mM activated sodium orthovanadate) and with protease inhibitor cocktail (Roche # 11873580001). Protein concentration in the lysates was determined by BCA assay (Thermo Fisher, #23227). Equal amounts of proteins were resolved in 14-6% Bis-Tris gels by gel electrophoresis at 110-150 V for 90 min using 1x Bis-Tris Running Buffer. The resolved proteins were transferred into Amersham™ Protran® nitrocellulose (0.45 µm, #15259794) membrane and blocked with Milk (5% in TBST). Membranes were then incubated overnight at 4°C with respective primary antibodies. On the following day, membranes were washed 3 times with TBST and incubated with respective secondary HRP antibodies for 1 h at room temperature. Membranes were then washed and signals were detected using ECL (Biorad, Clarity Western ECL Substrate, # 1705061) in the Bio-Rad Chemidoc imager as per the manufacturer’s protocol.

For tissue immunoblotting, respective snap frozen tissue samples were individually homogenized using Cellcrusher (Cellcrusher, Cork, Ireland) tissue pulveriser. The homogenized tissues were weighed and incubated in 10X volume of RIPA buffer (50 mM Tris–HCl pH 8, 150 mM NaCl, 1 mM EDTA, 1% NP-40, 1% Na-deoxycholate, 0.1% SDS) supplemented with protease inhibitor cocktail and phosphatase inhibitor cocktail for 30 mins on ice. At 15-minute intervals, tissue was pulverized for better extraction of proteins. Lysates were clarified by centrifugation at 14,000 rpm for 15 min at 4°C. Protein concentration was determined using the BCA assay (Thermo #23227). An equal amount of protein was resolved in a 6–14% Bis-Tris gel, and immunoblotting was performed as mentioned above

### 6. Microscopy-based *mito*-QC assay in cell lines

Cells stably expressing the *mito*-QC sensor were generated by retroviral transduction. The stable cells were seeded on a coverslip and incubated overnight in standard culture conditions. The following day, cells were treated with the respective drug as mentioned above for 48 hours. At the end of treatment, cells were washed three times with PBS and fixed at room temperature for 10 min in 3.7% Paraformaldehyde (PFA) buffered with 200m HEPES, pH 7.0. The fixed cells were washed and incubated for 15 min at RT with DMEM-HEPES (10mM, pH 7.0) containing 0.02% NaN3 to quench PFA. At the end of incubation, coverslips containing the cell were washed twice with PBS and mounted using Prolong Glass Antifade Mountant (Invitrogen, # P36984) and imaged on a Nikon Eclipse Ti2 widefield microscope with a 63X objective. Quantitation of the number of red-only puncta, indicating mitolysosomes, was performed using a semiautomated *mito*-QC counter in Image J software as described previously (*33*).

### 7. High Resolution Respirometry

FBXL4 KO cells stably expressing FLAG-empty vector or FLAG-FBXL4 were treated with either MK-8722 alone or in combination with BAY-3827 for 48 hours. Cells were then trypsinized and counted. 1×10^6^ cells were resuspended in 100 μl of Miro5 (Oroboros, #60101-01) mitochondrial respiration buffer. Samples were then loaded into the thermostated oxygraphic chamber at 37°C with continuous stirring (Oxygraph-2 k, Oroboros instruments, Innsbruck, Austria) to measure mitochondrial respiration. Cells were then permeabilized with digitonin (10 μg /10^6^ cells), and mitochondrial respiration for different respiration states was measured using the Substrate-Uncoupler-Inhibitor titration protocol number 8 (SUIT-008) as per the Oroboros protocol. Briefly, routine or leak respiration (PMLN) was first measured in the absence of any mitochondrial substrates and ATP. Mitochondrial respiration from Complex I (PGMp) was then measured by sequential injection of pyruvate (5 mM), Malate (2 mM), ADP (2.5 mM), and Glutamate (10 mM). Mitochondrial membrane integrity (PMcp) was tested by adding cytochrome c (10 μM) between the ADP and Glutamate injection steps. Any samples showing more than 10% increase in oxygen flux post cytochrome c addition was discarded as they indicate mitochondrial membrane damage. Combined contribution of Complex I and II (PGMSp, OXPHOS) on mitochondrial respiration was then measured with the injection of Succinate (10 mM). Uncoupled respiration (PGMSE, ETS) was next measured with CCCP titration. Residual mitochondrial respiration (ROX) was then measured by sequential addition of Rotenone (0.5 μM) and Antimycin (2.5 μM). The ROX oxygen flux value was subtracted from each of the above mitochondrial respiration states to quantify mitochondrial respiration of each state. Results are expressed in pmol/(s*cells).

### 8. Animal studies

All animal breeding and experimental procedures were approved by the University of Dundee Ethical Review Committee, the Named Veterinary Surgeon, and the Compliance Officer. All work was conducted under a UK Home Office project licence in accordance with the Animals (Scientific Procedures) Act 1986.

Two independent animal studies were performed. For both, mice were bred in-house and housed in sex-matched groups of 2–5 animals under a 12:12 h light–dark cycle at an environmental temperature of 21 ± 1 °C with 55–65% relative humidity. Food and water were provided *ad libitum*. Approximately 4-month-old male and female *mito*-QC mice on C57BL/6J background were used for all experiments (*40*).

#### Study 1: FBXL4 inhibition and AMPK activation

This study investigated the effects of FBXL4 inhibition (MLN-4924), AMPK activation (MK-8722), and combined treatment on mitophagy across multiple tissues. A total of 32 animals (n = 8 per group) were used, based on power calculations from our published in vivo data (*14*). Animals of both sexes were randomly assigned to groups and received once-daily intraperitoneal injections for 5 consecutive days. The vehicle for MLN-4924 consisted of 5% DMSO and 20% (w/v) β-cyclodextrin (Sigma #H5784) in saline, and it was administered at a dose of 50 mg/kg (MedChem Express #HY-70062), adapted from (*34*). MK-8722 was injected at a dose of 30 mg/kg based on (*35*). MK-8722 was formulated in 0.25% (w/v) methylcellulose, 5% (v/v) polysorbate 80, and 0.02% (w/v) sodium lauryl sulfate in sterile water.

#### Study 2: Metformin treatment

This study assessed the effects of metformin on hepatic mitophagy. A total of 20 animals (n = 10 per group) were included. Animals received once-daily intraperitoneal injections of either saline (vehicle control) or metformin at 150 mg/kg, dissolved in saline, for 5 consecutive days based on (*44*).

### 9. Blood glucose measurement

Blood glucose levels were assessed by blood micro-sampling in all animals at two time points: 30 min prior to and 60 min after the first intraperitoneal injection with vehicle or compounds mentioned above. For micro-sampling, the tail was punctured using a 25-gauge needle, and small blood droplets were collected directly onto glucose test strips. Glucose levels were quantified with a digital glucometer (CONTOUR® Meter).

### 10. Tissue collection and processing

Animals were anesthetized by intraperitoneal injection of 20% (v/v) pentobarbital (Euthatal, Merial) in PBS after 4 hours of the final intraperitoneal injection of respective compounds and vehicle. To eliminate blood from tissues, animals were transcardially perfused with PBS, after which various organs were collected and divided into two parts. One part was immediately fixed in 3.7% paraformaldehyde (PFA) prepared in 200 mM HEPES buffer (Sigma, #P6148; pH 7.0) overnight at 4 °C. The other portion was flash-frozen in liquid nitrogen for subsequent biochemical analyses by immunoblotting as mentioned above. Fixed tissues were washed three times with PBS containing 0.1% Azide and stored in 30% (w/v) sucrose in PBS for cryopreservation until use. Liver tissues were embedded in OCT compound (Agar Scientific, #AGR1180) and cryosectioned (10 µm) using a Leica CM1860UV cryostat. The sections were mounted on glass slides (Leica, #3800202) and air-dried in a fume hood. Sections were washed three times with PBS and counterstained with Hoechst 33258 (1 µg/ml, 5 min, room temperature). Excess stain was removed by two additional PBS washes, and slides were mounted with Vectashield Antifade Medium (Vector Laboratories, H-1000) and sealed with high-precision cover glasses (Marienfeld, #0107222) using transparent nail polish.

### 11. Immunolabeling of brain free-floating sections

Sagittal brain sections of 50 µm thickness were generated using a sledge microtome (Leica SM2010R), and slices were stored in PBS containing 0.1% Azide at 4°C until further processing. Free-floating sections were permeabilized three times for 5 min in DPBS (Gibco, #14190094) containing 0.3% Triton X-100 (Sigma Aldrich, #T8787). Samples were then incubated for 1 h in blocking buffer composed of DPBS supplemented with 10% goat serum (Sigma Aldrich, G9023) and 0.3% Triton X-100. The sections were then stained with anti-calbindin-D28k (Swant, #CB38, 1:1000 in blocking buffer) by overnight incubation of free-floating sections at room temperature. On the following day, sections were washed twice (8 min each) in DPBS with 0.3% Triton X-100 and incubated for 1 h in blocking buffer containing Pacific Blue anti-rabbit IgG secondary antibody (Invitrogen P10994, 1:200). After two additional 8-min washes in DPBS with 0.3% Triton X-100, sections were mounted on adhesive slides (Leica, #3800202) with coverslip using Vectashield Antifade Mounting Medium (Vector Laboratories, H-1000) and sealed with transparent nail polish.

### 12. Tissue confocal microscopy and quantitation of mitophagy in vivo

Tissue sections were imaged using an LSM 880 Airyscan microscope with Plan-Apochromat 63x/1.4 Oil DIC M27 objective. Approximately 10 images or 80% of the tissue area (for Hippocampus) were imaged across two to three sections for each tissue sample per individual animal. Quantification of the number of mitolysosomes was performed using Volocity® software (version 6.3, Perkin-Elmer) as described previously (*57*). In brief, all channels were processed using the software’s default fine filtering to reduce background noise. For liver analysis, EGFP fluorescence was thresholded to define the tissue area, while for brain sections, Pacific Blue fluorescence was thresholded to define the neuronal area. Mitolysosomes were identified by detecting peaks in the mCherry/EGFP ratio channel. The false-positive signals in areas of low reporter expression were excluded by applying an additional threshold for mCherry intensity. Threshold values were kept constant for all the tissues within a given experimental set. Mitophagy was quantified as the mean number of mitolysosomes per unit tissue area (µm²).

### 13. Statistical analysis

GraphPad Prism software (Version 10.1.0.316) was used for the statistical analysis. All data are represented as mean ± SEM. The number of biological replicates for each experiment is mentioned in the figure legends. Various statistical tests used to determine significance between groups are mentioned in the figure legend.

## ACKNOWLEDGEMENTS

I.G.G. was funded by the Medical Research Council, UK (MC_UU_00038/2) and a research alliance grant from the Novo Nordisk Foundation Center for Basic Metabolic Research (NNF CBMR) at the University of Copenhagen. NNF CBMR is an independent Research Center, partially funded by an unconditional donation from the Novo Nordisk Foundation (Grant number NNF18CC0034900 and NNF23SA0084103). RWT is funded by the Wellcome Centre for Mitochondrial Research (203105/Z/16/Z), LifeArc, the Medical Research Council (MR/W019027/1), the Lily Foundation, the UK NIHR Biomedical Research Centre for Ageing and Age-related disease award to the Newcastle upon Tyne Foundation Hospitals NHS Trust and the UK NHS Highly Specialised Service for Rare Mitochondrial Disorders of Adults and Children.

---

**Figure S1:**
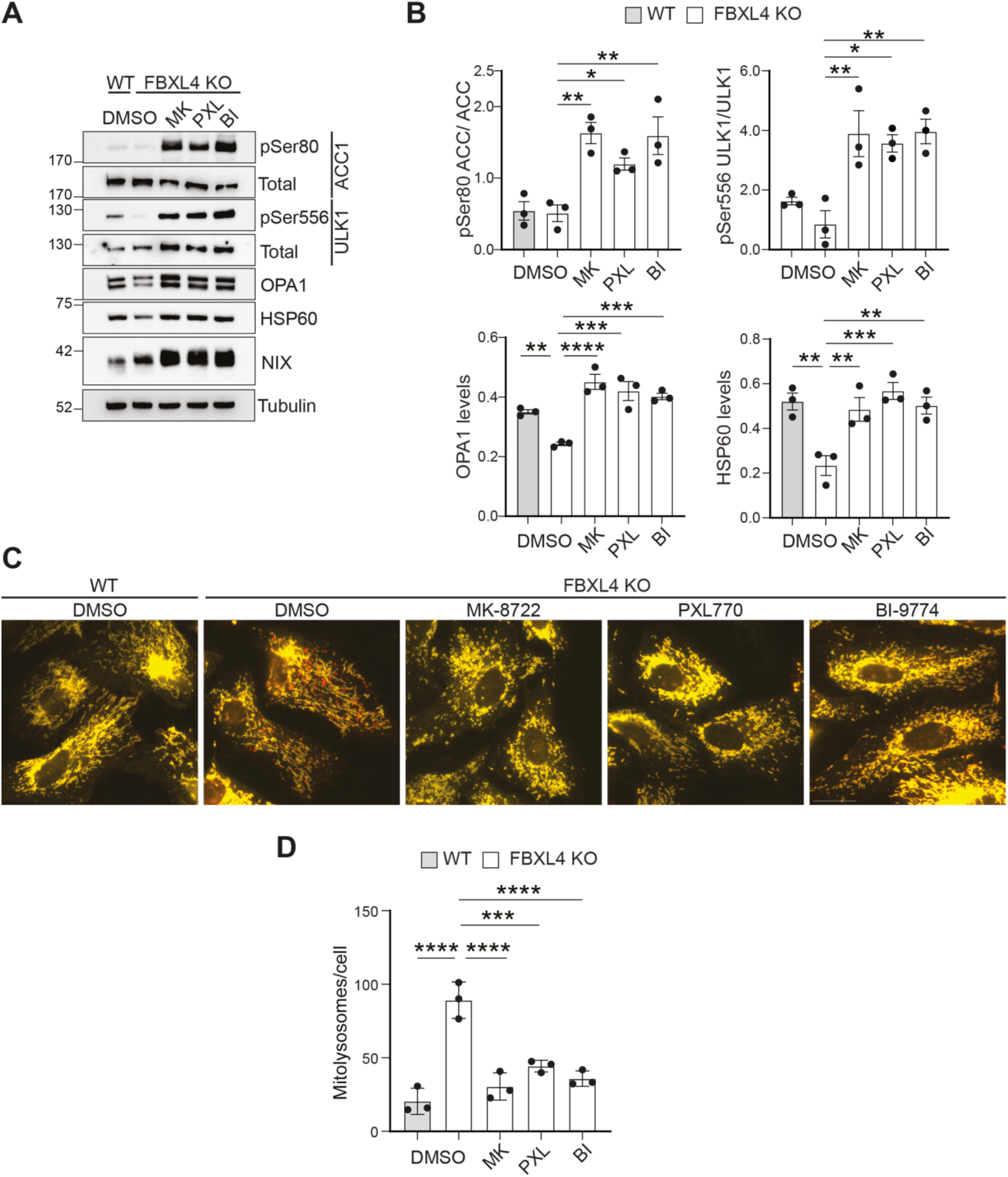
AMPK activators inhibit FBXL4 induced mitophagy. (**A**) Representative immunoblot of indicated proteins from ARPE-19 wildtype (WT) and FBXL4 knockout (FBXL4 KO) treated with/without MK8722 (MK, 10μM), PXl-770 (PXL, 50 μM) and BI-9774 (BI, 10μM) for 48 hours. (**B**) Quantification of the immunoblot data in panel A with protein levels normalised to loading controls. Data represents mean ± SEM (n=3), significance calculated using ordinary one-way ANOVA and Dunnett’s multiple comparisons test. (**C**) Representative wide-field images of WT and FBXL4 KO ARPE-19 *mito*-QC cells, treated as in A. Red puncta represent mitolysosomes while mitochondrial network appears yellow. Scale bar: 20 μm. (D) Mitolysosomes quantitation per cell. Data represents mean ± SEM (n=3), significance calculated using ordinary one-way ANOVA and Dunnett’s multiple comparison test. For all the experiments statistical significance is depicted as *p<0.05, **p<0.01, ***p<0.001 and ****p<0.0001.

**Figure S2:**
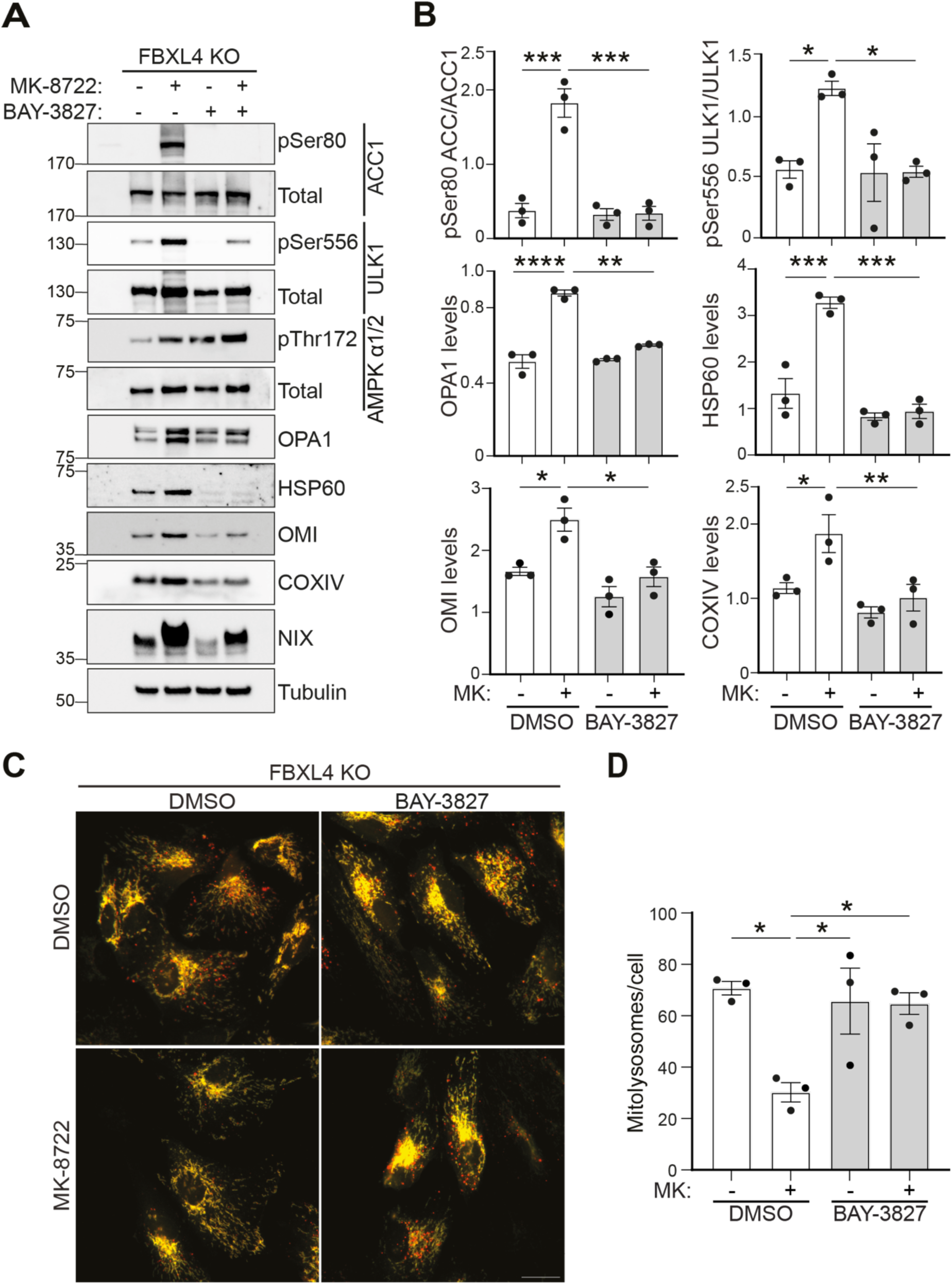
MK-8722 suppresses mitophagy in an AMPK-dependent manner. (**A**) Representative immunoblots of indicated proteins from FBXL4 knockout (FBXL4 KO) cells treated with MK-8722 (MK, 10 μM) and BAY-3827 (5 μM) for 48 hours. (**B**) Quantification of the immunoblot data in panel A with protein levels normalised to loading controls. Data represents mean ± SEM (n=3), significance calculated using ordinary one-way ANOVA and Tukey’s multiple comparisons test. (**C**) Representative wide-field images of WT and FBXL4 KO ARPE-19 *mito*-QC cells, treated with MK-8722 (MK, 10 μM) and BAY-3827 (5 μM) for 48 hours. Red puncta represent mitolysosomes while mitochondrial network appears yellow. Scale bar: 20 μm. (**D**) Quantification of the number of mitolysosomes per cell. Data represents mean ± SEM (n=3), significance calculated using ordinary one-way ANOVA and Tukey’s multiple comparison test. For all the experiments statistical significance is depicted as *p<0.05, **p<0.01, ***p<0.001 and ****p<0.0001.

**Figure S3:**
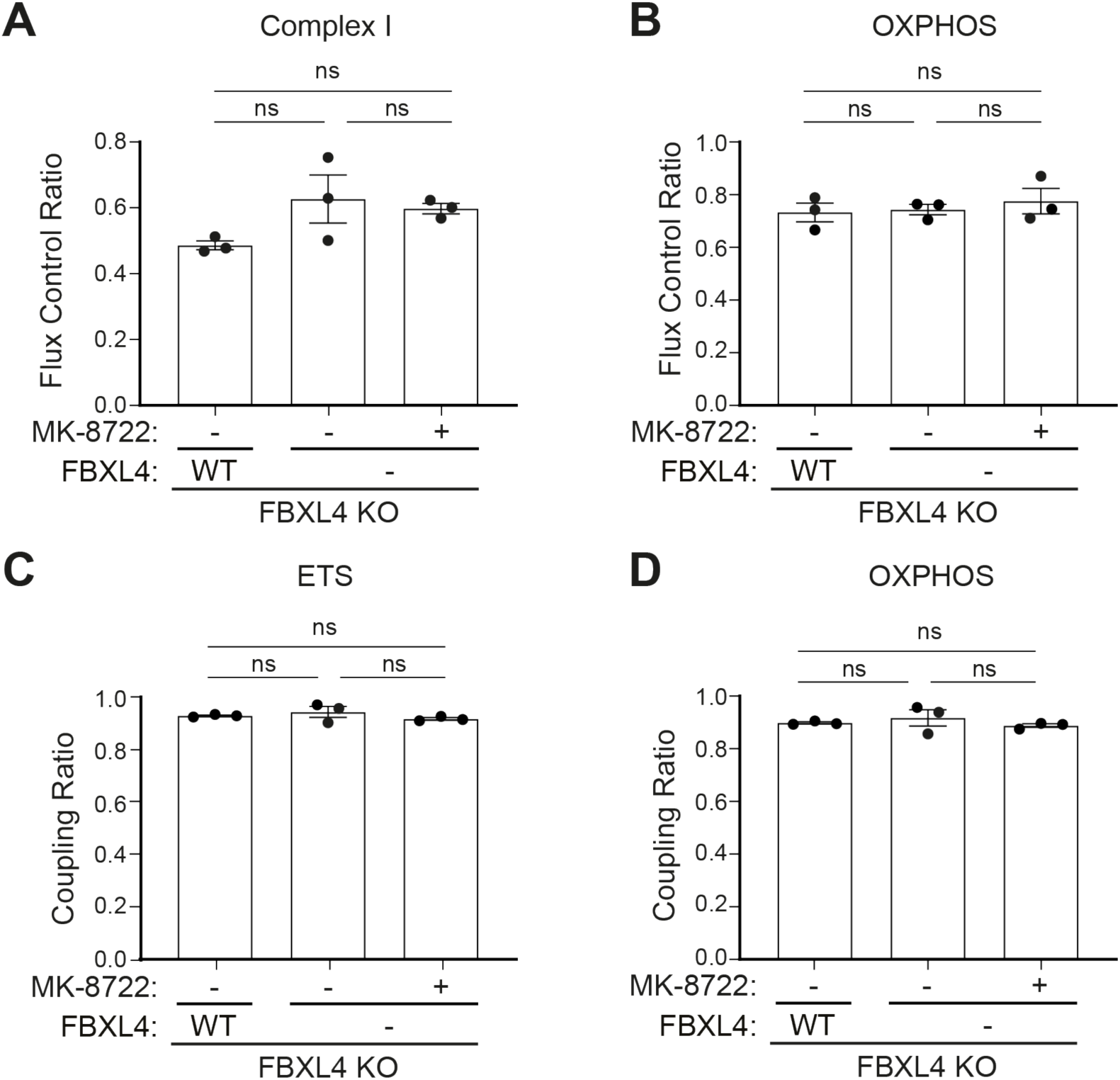
FBXL4 and AMPK do not alter respiratory capacity of mitochondrion. (**A-B**) Graph representing Flux control ratio (FCR) of Complex I (A) and OXPHOS (B) states measured from FBXL4 KO cells either expressing FLAG-empty vector or FLAG-FBXL4 WT treated with MK-8722 (10 μM) for 48 hours by high-resolution respirometry. FCR denotes the ratio of oxygen flux of Complex I and OXPHOS normalized to respective ETS state. Data represents mean ± SEM (n=3), significance calculated using ordinary one-way ANOVA and Tukey’s multiple comparison test. (**C-D**) Graph representing coupling ratio (CR) of ETS (C) and OXPHOS (D) states measured from FBXL4 knockout cells either expressing FLAG-empty or FLAG-FBXL4 WT treated with MK-8722 (10 μM) for 48 hours by high-resolution respirometry. CR denotes the ratio of net oxygen flux from ETS and OXPHOS normalized to respective ETS and OXPHOS states. For all the experiments ns depicts nonsignificant change.

**Figure S4:**
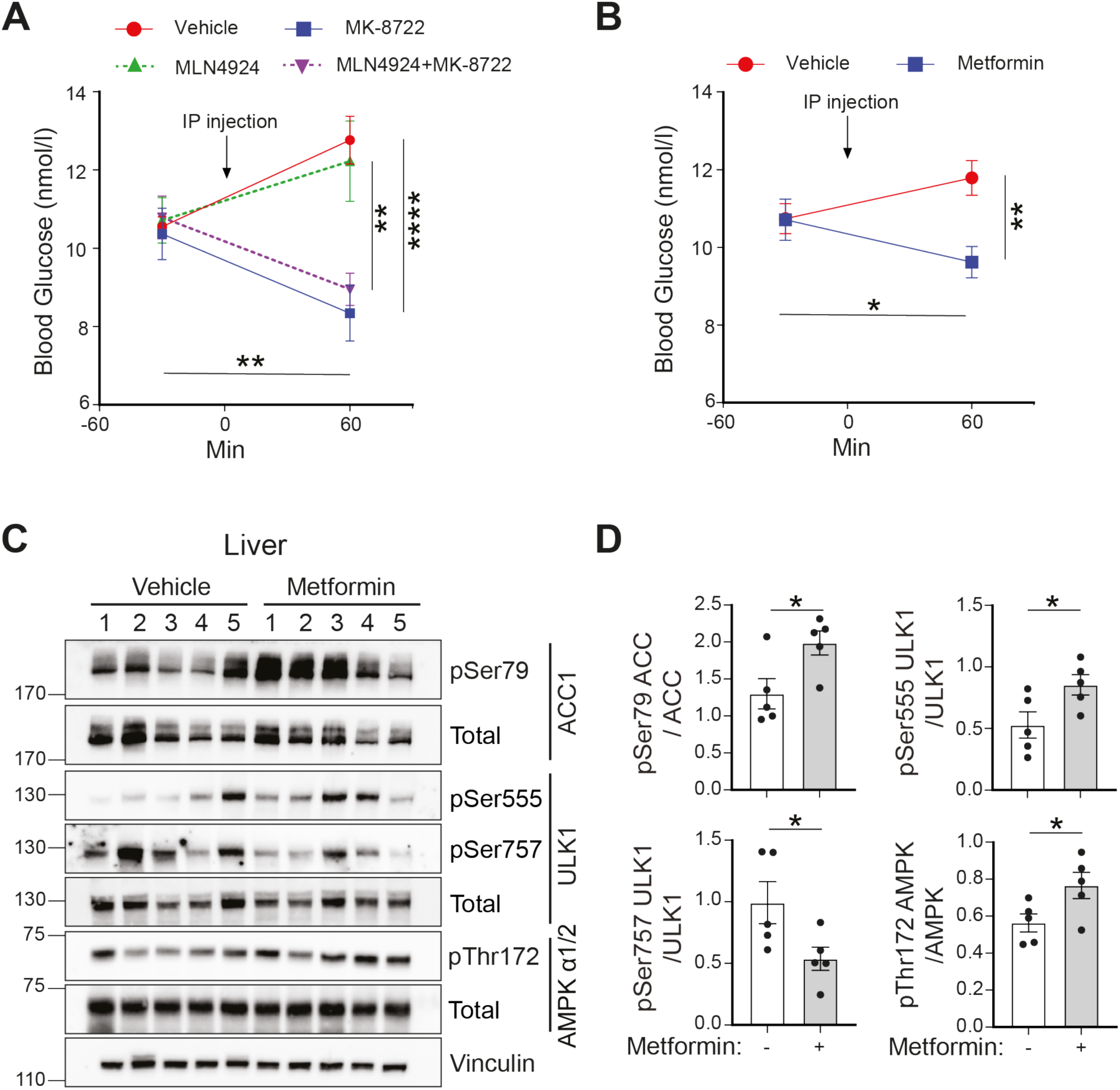
Effect of MK-8722 and Metformin on blood glucose levels and AMPK signalling. (**A**) Graph representing blood glucose level of *mito*-QC mice monitored 30 min pre- and 60 min post day 1 inter-peritoneal injection of vehicle, MK-8722 (30mg/kg), MLN-4924 (50mg/kg) by blood micro-sampling. Data represents mean ± SEM (n=8 mice), significance calculated using two-way ANOVA and Tukey’s multiple comparisons test. (**B**) Graph representing blood glucose level of *mito*-QC mice monitored 30 min pre- and 60 min post day 1 inter-peritoneal injection of vehicle or Metformin (150 mg/kg) by blood micro-sampling. Data represents mean ± SEM (n=10 mice), significance calculated using two-way ANOVA and uncorrected Fisher’s LSD test. (**C**) Representative immunoblots of indicated proteins from liver lysates from *mito*-QC mice injected with vehicle or metformin (150 mg/kg) for 5 days. (**D**) Quantification of the immunoblot data in panel C. Data represents mean ± SEM (n=5), significance calculated using unpaired t-test. For all the experiments statistical significance is depicted as *p<0.05, **p<0.01, ***p<0.001, ****p<0.0001.

**Figure S5:**
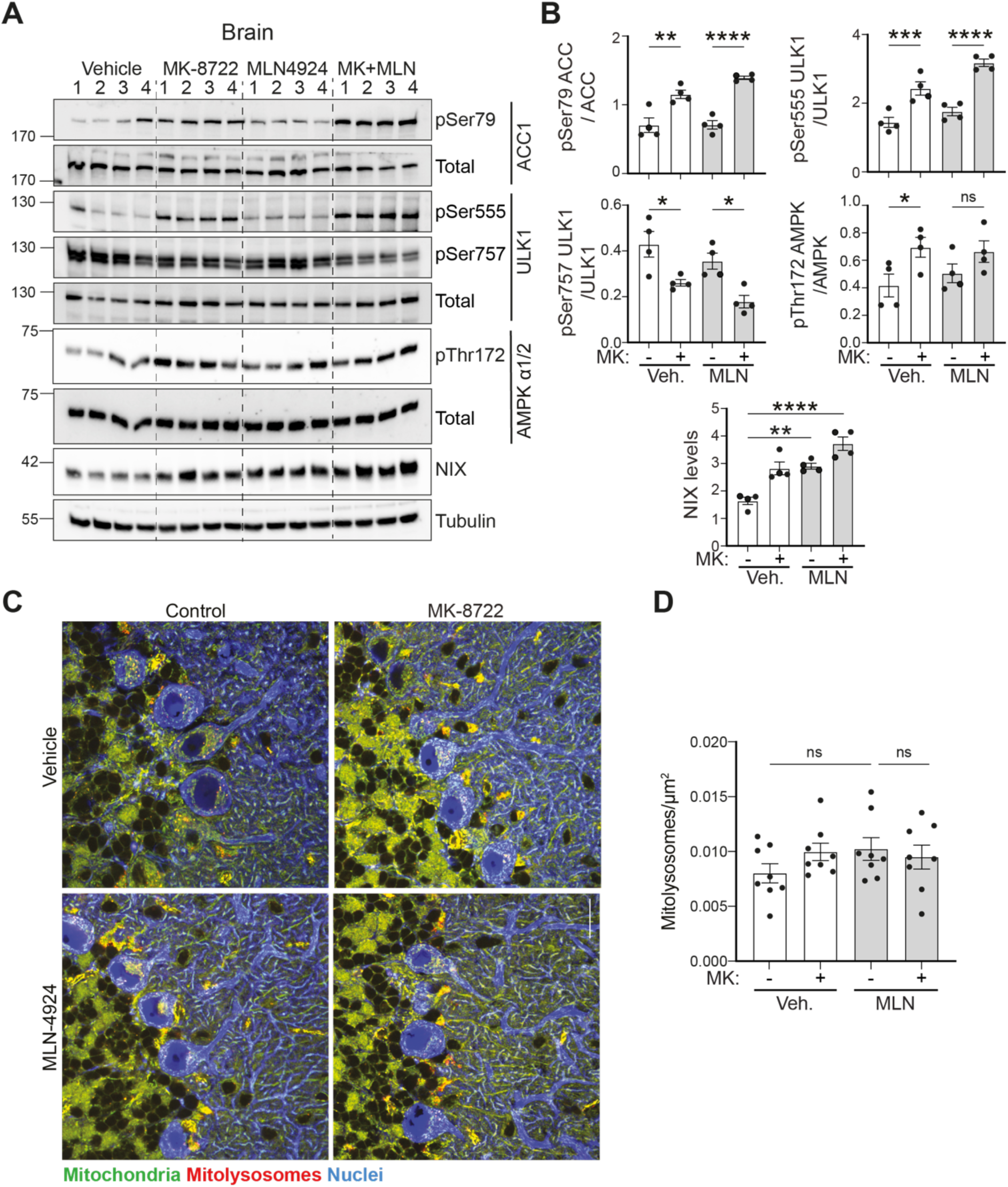
Effect of MK-8722 on AMPK signalling and mitophagy in the brain. (**A**) Representative immunoblots of indicated proteins from whole brain lysates from *mito*-QC mice dosed with vehicle, MK-8722 (30 mg/kg) and MLN-4924 (MLN, 50 mg/kg) for 5 days. Each lane represents an individual mouse. (**B**) Quantification of the immunoblot data in panel A with protein levels normalised to loading controls. Data represents mean ± SEM (n=4 mice), significance calculated using ordinary one-way ANOVA and Sidak’s multiple comparisons test. (**C**) Representative confocal micrographs of Calbindin immunolabelled Purkinje neurons (Blue) in the Cerebellum from the brain sections of *mito*-QC mice dosed with vehicle, MK-8722 (30 mg/kg) and MLN-4924 (MLN, 50 mg/kg) for 5 days. Red puncta indicate mitolysosomes, whereas the mitochondrial network is depicted in green. Scale bar: 20 μm. (**D**) Quantification of the number of mitolysosomes normalized to calbindin labelled Purkinje neuron area. Data represents mean ± SEM (n=8), significance calculated using ordinary one-way ANOVA and Sidak’s multiple comparisons test. For all the experiments statistical significance is depicted as *p<0.05, **p<0.01, ***p<0.001, ****p<0.0001 and ns: nonsignificant.

